# Tumour genotype shapes blood biomarker expression for use in pancreatic cancer detection and diagnosis

**DOI:** 10.1101/2025.08.15.670486

**Authors:** Marta Canel, David W Lonergan, Cameron Ferguson, Philippe Gautier, Jennifer P Morton, Alex von Kriegsheim, Alan Serrels

## Abstract

Typically diagnosed late, when systemic metastasis has already occurred, pancreatic ductal adenocarcinoma (PDAC) has one of the worst 5-year survival rates of any cancer type. For many patients with advanced disease, current chemotherapy regimens offer only modest benefit despite significant toxicity and surgical resection, the only treatment option with curative potential, is not possible. Therefore, while new treatments are much needed, diagnosing patients at an ‘earlier’ disease stage when surgery remains possible and the window of opportunity for treatment response is longer will be critical to improving patient outcomes. In this regard, the identification of biomarkers from biospecimens that can be easily sampled from patients remains the focus of considerable research, however success has not been forthcoming. Using a suite of novel genetically defined murine isogenic models of early PDAC, engineered using CRISPR-Cas9 gene editing, we sought to address whether loss-of-function mutations in common driver genes, and thus the genetic heterogeneity inherent to the disease, may represent an important confounding factor in the identification of a one-size-fits-all biomarker suitable for early detection. Focussing on the multi-omics analysis of blood, we show that both loss of *Cdkn2a* and / or *Smad4* on the background of a *Kras^G12D^ Trp53^-/^*^-^ genotype has profound effects on the profile of differentially expressed RNA species including protein coding RNAs, lncRNAs, snoRNAs, scRNAs, snRNAs and miRNAs, and on plasma protein expression, when compared to both healthy controls and chemically induced pancreatitis. In addition, we find that loss of *Smad4*, a genomic event that occurs following progression from PanIN to PDAC, substantially limits the availability of blood biomarkers. These findings identify the need to move towards genotype-specific biomarker signatures and uncover a potential role for *Smad4* loss in limiting opportunities for the early detection of pancreatic cancer.

## INTRODUCTION

Often asymptomatic during early development and lacking effective screening tools to facilitate early diagnosis, Pancreatic Ductal Adenocarcinoma (PDAC) is most commonly diagnosed at an advanced disease stage (80-85% of patients) by which time treatment options are limited to palliative chemotherapy (1). With increasing incidence and little improvement in treatment options over the last 40 years, pancreatic cancer is on course to become the second leading cause of cancer death, only surpassed by lung cancer (2, 3). The overall 5-year survival rate for PDAC is ∼8%. However, for the minority of patients eligible for surgery, successful pancreatic resection can increase median 5-year survival to as much as 30% (Pancreatic Cancer Action Network). Therefore, while new treatments are much needed, it is clear that diagnosing patients at an earlier disease stage when surgery remains possible, and the window of opportunity for treatment is longer, will be critical to improving patient outcome (4).

PDAC has a relatively low incidence, with approximately 17 individuals for every 100,000 of the European population being diagnosed every year (Cancer Research UK (CRUK) statistics). As a result, General Practitioners (GPs) in the United Kingdom will likely only see on average one new pancreatic cancer patient every 5 years. Many of the symptoms that occur during the development of the disease, including abdominal pain, mid-back pain, loss of appetite, indigestion, unexplained weight loss, fatigue etc, are vague and easily confused with more common health problems, often contributing to delayed diagnosis or misdiagnosis, the latter of which occurs in >30% of cases. As a result, nearly half of all pancreatic cancer patients are diagnosed after becoming unwell and requiring a visit to Accident and Emergency (Pancreatic Cancer UK (PCUK) and CRUK) hospital care. Diagnosis downstream of a GP referral can improve 1-year survival by up to three-fold (PCUK), highlighting the need to implement diagnostic tests that can aid GPs in identification of patients requiring urgent referral for further investigation. Hence, there is considerable interest in the identification of potential biomarkers to aid earlier diagnosis, helping primary care physicians triage patients most in need of urgent care.

Currently, carbohydrate 19-9 (CA19.9) is the only biomarker widely used in the diagnosis of pancreatic cancer. However, it is more effective in prognosis following post-operative recurrence than as a diagnostic marker for early disease (5, 6). Analysis of pancreatic tumours and various biofluids from patients has yielded a broad range of candidate biomarkers. However, none have ultimately shown the sensitivity and specificity required for adoption into clinical practice (7–9). We therefore sought to identify key factors that may confound the current strategy for biomarker discovery, which assumes that a one-size-fits-all biomarker or biomarker signature will be sufficient to support early detection.

Pancreatic cancer is a genetically heterogeneous disease. Extensive genomic profiling has shown that KRAS mutations have an extremely high prevalence in PDAC (>90%) with a variety of mutations in codon-12 e.g. KRAS^G12D^ (∼50%), KRAS^G12V^ (∼30%), KRAS^G12R^ (∼15%), accounting for over 90% of KRAS mutations in pancreatic cancer (10). However, KRAS mutation alone is not sufficient to drive pancreatic cancer development and additional loss-of-function mutations in one or more tumour suppressor genes, most commonly *TP53* (∼74%), *CDKN2A* (∼44%) and *SMAD4* (∼22%), are required to promote disease development and progression, resulting in a range of genetic backgrounds across the patient population (11, 12). We therefore set out to determine whether genetic heterogeneity may represent a key confounding variable in the identification and validation of robust biomarkers for the early detection of PDAC and, in doing so, identify whether a step-change towards genotype-specific biomarker signatures is required.

Using the multi-omics analysis of blood, together with a novel suite of isogenic cell-based models of PDAC generated using CRISPR-Cas9 genome editing of primary acinar cells isolated from the pancreas of C57BL/6 mice, we find that loss of function of *Cdkn2a* and / or *Smad4* fundamentally reprograms RNA and protein biomarker expression on a *Kras^G12D^ Trp53^-/-^* background. Comparison of tumour bearing mice to age-matched healthy controls followed by filtering for RNAs and proteins also regulated as a consequence of pancreatitis, revealed almost no biomarkers common across *Kras^G12D^ Trp53^-/-^*, *Kras^G12D^ Trp53^-/-^ Cdkn2a^-/-^*, *Kras^G12D^ Trp53^-/-^ Smad4^-/-^* and *Kras^G12D^ Trp53^-/-^ Cdkn2a^-/-^ Smad4^-/-^* genotypes. Thus, we identify tumour genetic background as a major confounding factor in biomarker discovery and pan-PDAC validation. Surprisingly, we also find loss of *Smad4*, either alone or in combination with *Cdkn2a*, to be a deleterious event, severely constricting the RNA biomarker pool, thereby limiting opportunities for biomarker discovery. Lastly, comparison of enriched RNAs in matched tumour and blood samples revealed a low level of concordance, implying that PDAC tumours do not represent a good surrogate for identification of blood-based biomarkers. These data, generated using a genetically defined model system, provide rationale underpinning the failure of current candidate biomarkers and strongly support the need to incorporate genomic information into biomarker discovery efforts in the clinic, with a view to developing genotype-specific biomarker signatures that support both diagnosis and potentially also patient stratification.

## RESULTS

### Differential driver gene expression results in distinct cell states

To generate a disease relevant, genetically defined, model system with which to investigate the impact of common driver gene loss-of-function mutations on blood biomarker expression for the early detection of PDAC, we isolated primary acinar cells from the pancreas of a healthy adult C57BL/6 mouse and used CRISPR-Cas9 genome editing to engineer four common genotypes found in human PDAC: (1) *Kras^G12D^ Trp53^-/^*^-^, (2) *Kras^G12D^ Trp53^-/-^ Cdkn2a^-/-^*, (3) *Kras^G12D^ Trp53^-/-^ Smad4^-/-^* and (4) *Kras^G12D^ Trp53^-/-^ Cdkn2a^-/-^ Smad4^-/-^* (**Figure 1A** and **1B**). Briefly, murine acinar cells were engineered to express oncogenic *Kras^G12D^*using targeted homology-directed repair (HDR) CRISPR-Cas9 of the endogenous *Kras* gene and *Trp53* expression was simultaneously deleted using recombinant CRISPR-Cas9 gene editing. Following isolation of a clonal cell population positive for the expression of both KRAS^G12D^ and the ductal lineage marker keratin-19, and negative for the expression of p53, targeted CRISPR-Cas9 genome editing was used to delete p16INK4A and p19ARF (both protein products of the *Cdkn2a* gene), and / or SMAD4 expression. To broadly characterise the impact of common driver genes on cell signalling we next analysed whole cell protein lysates isolated from cells in culture using Reverse Phase Protein Arrays (RPPA) (**Figure 1C**). This analysis identified the differential expression / activation of distinct intracellular signalling pathways dependent on genotype, implying that the combination of driver genes present results in distinct cell states. We next implanted 1 x 10^5^ cells of each cell line into the pancreas of C57BL/6 mice and culled mice 3-weeks post implantation to determine the effects of different genotypes on primary tumour growth. We found tumour weights to be relatively modest at this time point, with *Kras^G12D^ Trp53^-/-^* tumours only weighing an average of 71 mg. In agreement with previously published observations (13–17), both *Cdkn2a* and / or *Smad4* loss was observed to promote primary tumour growth resulting in tumours weighing an average of 146 – 197 mg (**Figure 1D**). Given that tumours at this time point were relatively small and that biomarkers suitable for the early detection of PDAC should ideally be disease stage agnostic, such that time of sampling is irrelevant, we decided to focus on this time point for the analysis of blood to address the impact of tumour genotype on the expression of candidate circulating biomarkers.

**Figure 1.**
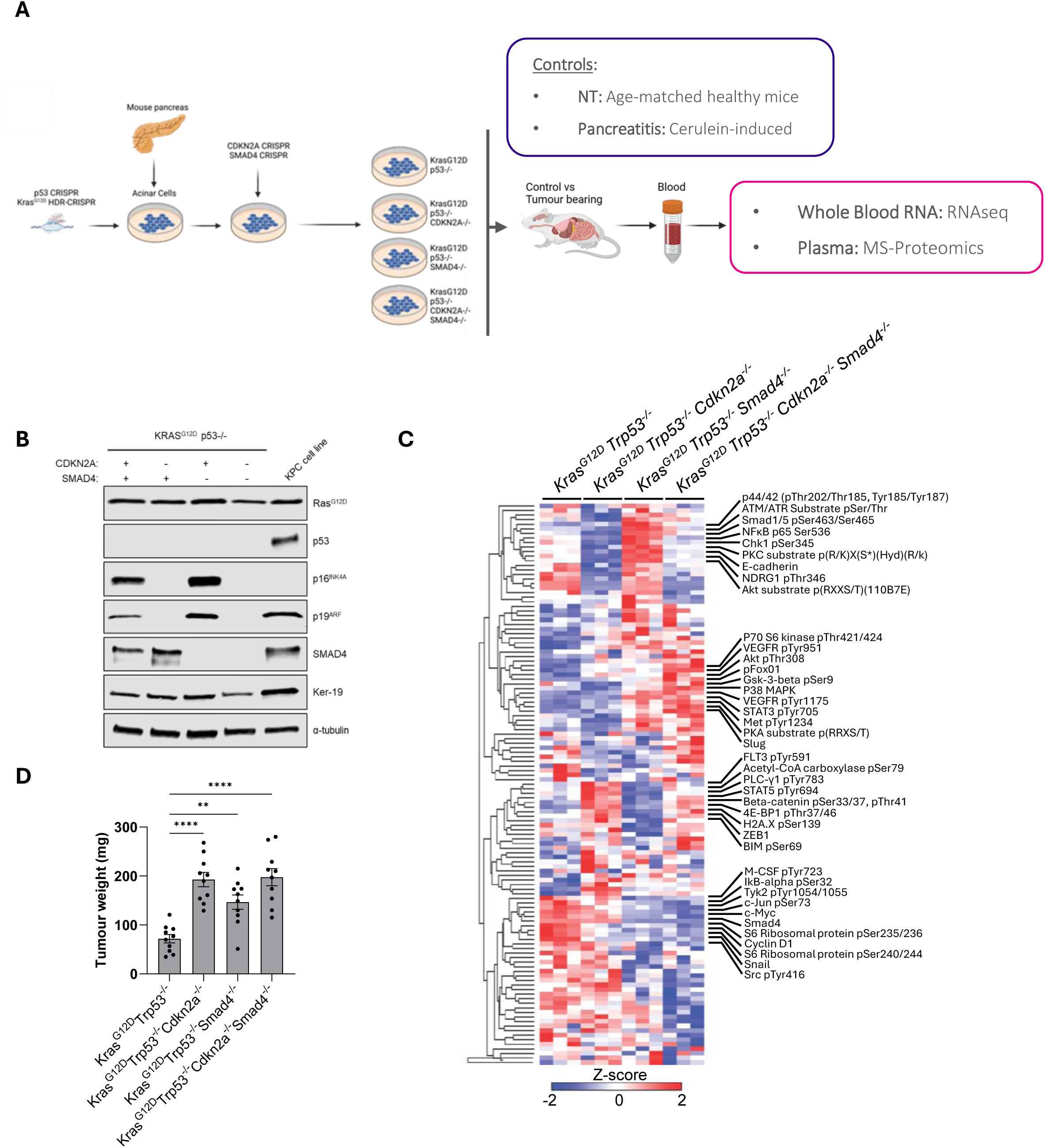
CRISPR-Cas9 genome editing of primary acinar cells to generate oncogenic cell-based models of PDAC representing different driver gene expression states. (A) Schematic describing the generation of cell-based PDAC models using CRISPR-Cas9 and subsequent experimental setup. (B) Representative western blot of whole cell lysates showing expression of KRAS^G12D^, p53, p16INK4A, p19ARF, SMAD4 and keratin-19. α-tubulin used as a loading control – image provided is representative of loading across all blots. (C) Reverse phase protein array analysis of whole cell protein lysates from cells in culture. Data Z-transformed and subject to hierarchical clustering. (D) Tumour weight 3-weeks post-implantation of 1×105 cells into the pancreas of C57BL/6 mice. N = 10 tumours per group. Data represented as mean ± s.e.m. Significance determined using ordinary one-way ANOVA with Tukey’s multiple comparison. **** p ≤ 0.0001, ** p ≤ 0.01

### Tumour genotype impacts circulating RNA biomarker expression

Blood-based biomarkers represent an attractive option for the early detection of PDAC, with a variety of RNA, DNA and protein species present. Therefore, we first isolated total RNA from blood 3-weeks post-implantation of either *Kras^G12D^ Trp53^-/^*^-^, *Kras^G12D^ Trp53^-/-^ Cdkn2a^-/-^*, *Kras^G12D^ Trp53^-/-^ Smad4^-/-^*or *Kras^G12D^ Trp53^-/-^ Cdkn2a^-/-^ Smad4^-/-^* cells into the pancreas of C57BL/6 mice. To enable identification of tumour-specific biomarkers we also isolated blood from age-matched healthy control animals and mice treated with cerulein to induce acute pancreatitis. To broadly assay the effects of tumour genotype on the circulating expression levels of different RNA species, we prepared sequencing libraries from total RNA following globin and ribosomal depletion. cDNA was sequenced using an Illumina NextSeq2000 and analysed as described in materials and methods. Principle component analysis (PCA) of biological replicates across the six experimental groups identified limited intra-group variability, except for the pancreatitis samples which appeared to cluster broadly into two groups (**Figure 2A**). To understand whether these two groups represented distinct biological effects of cerulein treatment, we identified differentially expressed RNAs (padj ≤ 0.05, log2fc ≥ 1 or ≤ -1) by pairwise comparison of each pancreatitis group to age-matched healthy controls (NT = no tumour). Cell type overrepresentation analysis based on genes significantly upregulated in pancreatitis GP1 and GP2 identified a predominant immune cell signature in GP1, while GP2 was enriched with a hepatocyte cell signature indicative of effects on the liver (**Supplementary Figure 1A**). Similar overrepresentation analysis based on Gene Ontology (GO) Biological Processes further confirmed the immune signature associated with GP1, while in contrast GP2 was dominated with GO terms associated with metabolic function (**Supplementary Figure 1B**). Therefore, we divided the pancreatitis samples into two groups: GP1 and GP2 based on the apparently distinct biological effects of the treatment. To first understand the effect of tumour genotype on the expression of candidate RNA biomarkers, we next identified differentially expressed RNAs (padj ≤ 0.05, log2 fc ≥ 1 ≤ -1) for each tumour genotype relative to NT controls (**Figure 2B**). This analysis identified only 31 upregulated RNAs and 2 downregulated RNAs to be conserved across the blood of all tumour bearing mice out of a total of approximately 25,000 RNAs identified. These findings imply that the pool of differentially expressed RNAs from which to develop a potential one-size-fits-all test is extremely limited. In contrast, we identified 314 upregulated and 19 downregulated RNAs unique to the *Kras^G12D^ Trp53^-/-^* genotype, 93 upregulated and 83 downregulated RNAs unique to the *Kras^G12D^ Trp53^-/-^ Cdkn2a^-/-^* genotype, and an additional 119 upregulated and 11 downregulated RNAs common between these two genotypes only. Unexpectedly, loss of *Smad4* had profound effects on the expression of potential RNA biomarkers, with only 1 upregulated and 1 downregulated RNA unique to the *Kras^G12D^ Trp53^-/-^ Smad4^-/-^* genotype, and 11 upregulated and 22 downregulated RNAs unique to the *Kras^G12D^ Trp53^-/-^ Cdkn2a^-/-^ Smad4^-/-^* genotype.

**Figure 2.**
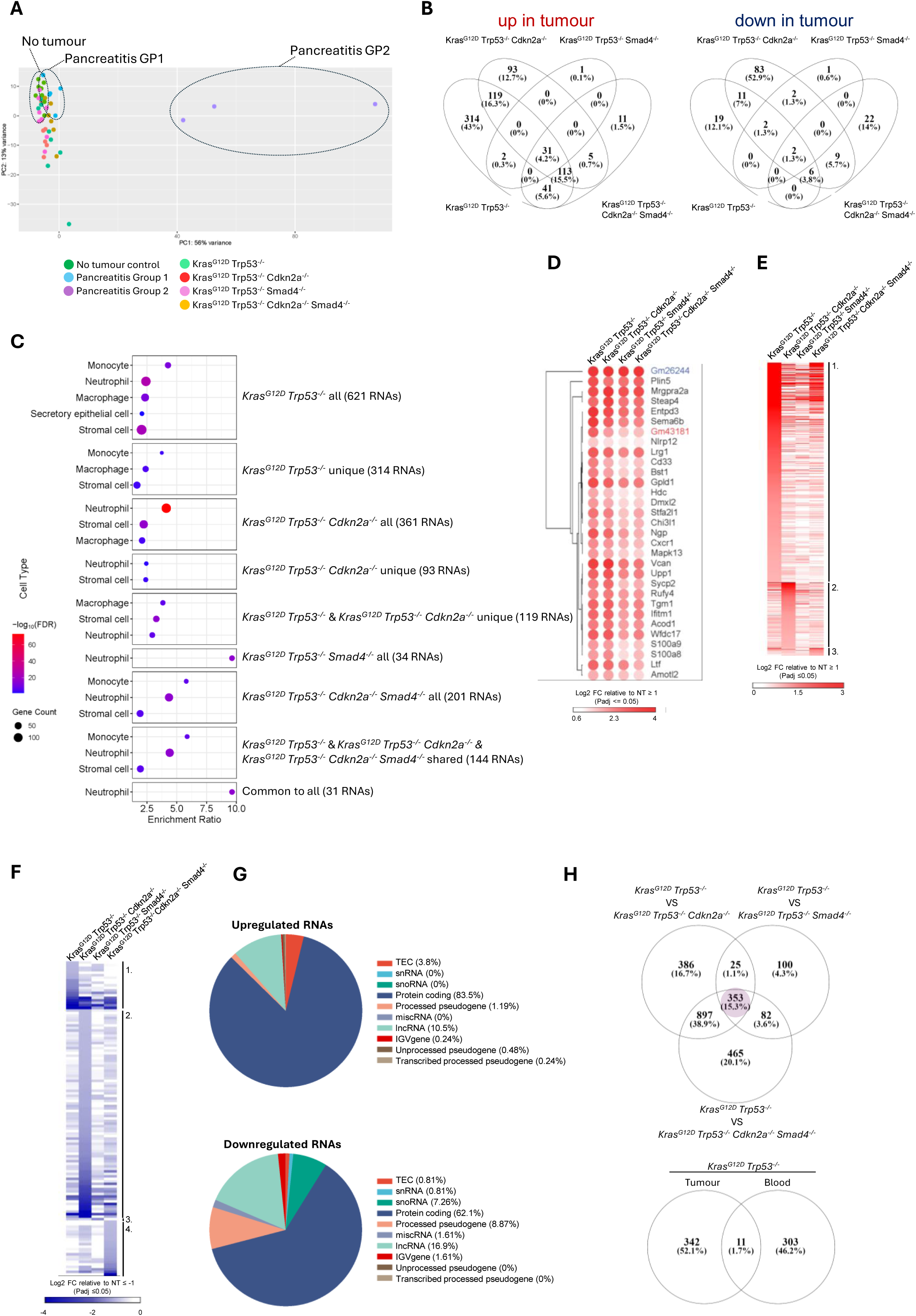
Tumour genotype impacts circulating RNA biomarker expression. (A) Principal Component Analysis of RNA sequencing data from biological replicates across all six experimental conditions. N = 8 biological replicates / group. (B) Venn diagram showing intersection between lists of differentially expressed RNAs identified following pairwise comparison of each tumour group to NT control. (C) Cell type overrepresentation analysis based on significantly upregulated RNAs present within different regions of intersection in B. (D) Hierarchical clustering of log2-fold change for the 31 RNAs in B upregulated across all four tumour genotypes relative to NT control. (E) Hierarchical clustering of significantly upregulated RNAs unique to each tumour group in B. (F) Hierarchical clustering of significantly downregulated RNAs unique to each tumour group in B. (G) Pie charts showing the percentage of different RNA species represented within the significantly up- and downregulated RNAs identified following pairwise comparison of all tumour groups to NT control. (H) *Top* – Venn diagram showing intersection between gene lists from pairwise comparison of *Kras^G12D^ Trp53^-/-^* tumour RNAseq data with all other tumour genotypes. 353 genes highlighted in pink are upregulated in *Kras^G12D^ Trp53^-/-^* tumours when compared to all other tumour genotypes. *Bottom* - Comparison of significantly upregulated RNAs in matched tumour and blood on a *Kras^G12D^ Trp53^-/-^* tumour background when compared to all other tumour genotypes.

To gain insight into the cellular source of the RNAs identified for each tumour group and what might underlie the differences observed, we next performed cell type overrepresentation analysis based on significantly upregulated RNAs (**Figure 2C**). Overrepresentation of a neutrophil signature was common to all groups and was the only enriched cell type signature identified from the 31 RNAs common to all tumour groups. However, an enriched neutrophil signature was also present in the 93 RNAs unique to the *Kras^G12D^ Trp53^-/-^ Cdkn2a^-/-^* tumour group, implying that tumour genotype may alter circulating immune cell state to reshape the available biomarker pool. Notably, stromal cell signatures were also present in unique subsets of RNAs from multiple tumour groups, implying that circulating stromal cell state may also be dependent on tumour genotype. While neutrophil and stromal cell signatures were common across the majority of tumour groups, an enriched macrophage signature was only present in *Kras^G12D^ Trp53^-/-^* and *Kras^G12D^ Trp53^-/-^ Cdkn2a^-/-^*tumour groups. Therefore, in addition to altered cell states, cell type abundance i.e. the composition of the circulating cellular milieu, is also potentially dependent on tumour genotype.

Given that many studies to date have attempted to identify biomarkers suitable for detection of pancreatic cancer irrespective of tumour genotype (8), we next asked whether the 31 RNAs upregulated across the four genotypes were differentially expressed at similar levels when compared to NT controls. While *Kras^G12D^ Trp53^-/^*^-^ and *Kras^G12D^ Trp53^-/-^ Cdkn2a^-/-^* tumour bearing mice showed similar patterns of expression, *Smad4* loss alone or in combination with *Cdkn2a* loss showed a trend towards a reduction in the log2 fold-change relative to NT (**Figure 2D**, **Supplementary Figure 2**), suggesting that loss of *Smad4* may negatively impact signal to noise, further limiting the number of RNAs with which to develop a robust pan-pancreatic cancer RNA signature. In contrast, many more differentially expressed RNAs were unique to a specific tumour group (**Figures 2E** and **2F**), implying that genotype-specific RNA signatures may offer a more fruitful approach for diagnosis.

Having sequenced total RNA to understand the relationship between tumour genotype and biomarker expression, our sequencing libraries were not specific to protein coding mRNAs. We therefore analysed the proportion of different RNA species present in the lists of differentially expressed RNAs unique to a given tumour group (**Figures 2E** and **2F**). While the majority of upregulated RNAs were protein coding (83.5%), over 10% of those identified were lncRNAs (**Figure 2G**). Notably, a smaller proportion of downregulated RNAs were protein coding (62.1%), with a greater frequency of lncRNAs, snoRNAs and processed pseudogenes. Therefore, the expression of both coding and non-coding RNAs is dependent on tumour genotype, and both represent potential biomarkers for diagnosis.

Many studies have set out to identify potential diagnostic biomarkers using transcriptomic datasets derived from PDAC tumour tissue (18). However, a diagnostic test to detect PDAC early would rely on sampling readily available biofluids as opposed to pancreatic tissue directly. To determine the conservation between tumour-specific RNAs present in matched tumour tissue and blood, we first used bulk tumour RNA sequencing data to identify differentially expressed RNAs upregulated specifically in *Kras^G12D^ Trp53^-/-^* tumours when compared with *Kras^G12D^ Trp53^-/-^ Cdkn2a^-/-^*, *Kras^G12D^ Trp53^-/-^ Smad4^-/-^* and *Kras^G12D^ Trp53^-/-^ Cdkn2a^-/-^ Smad4^-/-^* genotypes (**Figure 2H** *-top*). A similar analysis was performed using RNA sequencing data from total RNA isolated from matched blood. Direct comparison of upregulated RNAs from tumour and blood identified only 1.7% overlap (**Figure 2H** *-bottom*), suggesting that the tumour does not represent a reliable surrogate for identification of RNA blood biomarkers.

A key property of a robust biomarker for the detection of cancer is its specificity for the disease. For example, a good biomarker should not be differentially expressed in both pancreatic cancer patients and patients with unrelated pathologies of the pancreas. Pancreatitis, inflammation of the pancreas, is an acute or chronic condition that can present with symptoms overlapping with those of pancreatic cancer (19). Cancer itself is commonly associated with an inflammatory response. Therefore, we sought to understand the extent to which pancreatitis may further impact the pool of potential biomarkers available for cancer detection across the range of genotypes. Administration of cerulein, a 10-amino acid oligopeptide, is the most commonly used experimental approach to promote pancreatitis in mouse models (20, 21). Therefore, we used this approach to characterise changes in RNA expression in the blood in response to pancreatic inflammation. Pancreatitis samples were split into two groups (GP1 and GP2) based on PCA analysis (**Figure 2A** and **Supplementary Figure 1**) and pairwise comparison to each tumour genotype used to identify differentially expressed RNAs. The lists of differentially expressed RNAs relative to NT, GP1 and GP2 were then cross compared using Venn diagrams, with the point of intersection between the three RNA lists representing tumour-specific biomarkers for each tumour group (**Figures 3A** – **3D**). The incorporation of RNA signatures associated with pancreatic inflammation into this analysis resulted in a substantial reduction in the number of potential tumour-specific biomarkers, suggesting that many of the differentially expressed RNAs identified between tumour groups and NT controls were the consequence of more generalised pancreatic inflammation. Notably, only 87 RNAs remained as potential biomarkers for *Kras^G12D^ Trp53^-/-^* tumours, while only 45 RNAs remained as potential candidates for *Kras^G12D^ Trp53^-/-^ Cdkn2a^-/-^* tumours. Only 5 RNAs remained as potential biomarkers associated with *Kras^G12D^ Trp53^-/-^ Smad4^-/-^* tumours and no candidate tumour-specific RNAs remained for *Kras^G12D^ Trp53^-/-^ Cdkn2a^-/-^ Smad4^-/-^* tumours, as these were indistinguishable from pancreatitis GP1. Further comparison of the remaining tumour-specific RNAs between groups (**Figure 3E**) identified only 1 upregulated RNA and 1 downregulated RNA common to *Kras^G12D^ Trp53^-/-^, Kras^G12D^ Trp53^-/-^ Cdkn2a^-/-^* and *Kras^G12D^ Trp53^-/-^ Smad4^-/-^* genotypes. Rather, the majority of differentially expressed RNAs were specific to a given genotype (**Figures 3F** and **3G**). These findings not only support the need to move towards genotype-specific RNA-based biomarker signatures but also suggest that loss of *Smad4* may represent a major confounding factor in the identification and development of RNA-based diagnostic tests for pancreatic cancer detection. Interestingly, analysis of RNA species present in the remaining lists of upregulated and downregulated RNAs identified that protein coding RNAs were by far the most common to be upregulated (**Figure 3H**). However, snoRNAs were by far the most common downregulated RNA type, further highlighting the potential utility of non-coding RNAs in the development of diagnostic tests.

**Figure 3.**
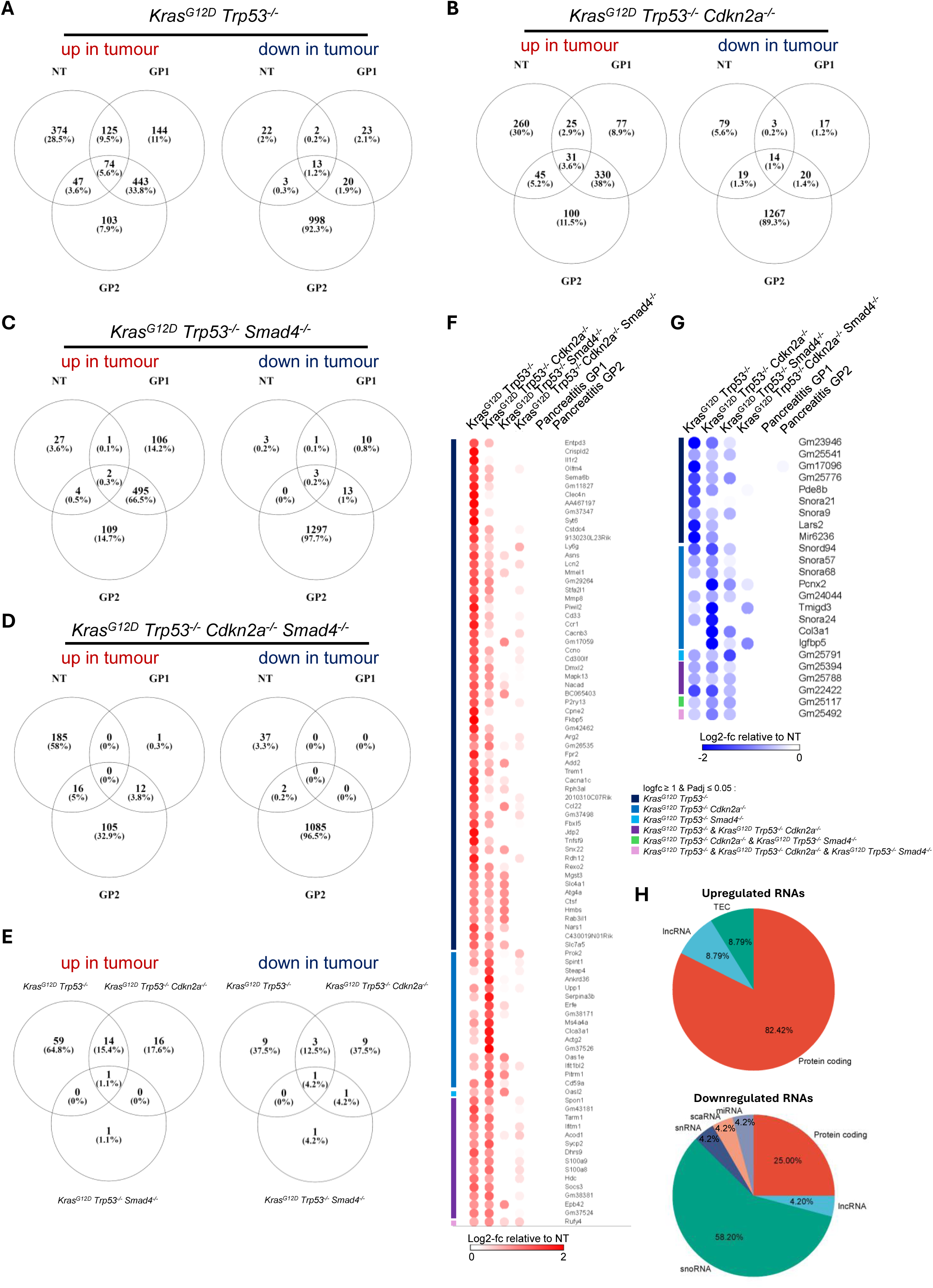
*Smad4* loss drives a blood RNA signature that is indistinguishable from pancreatitis. (A - D) Venn diagram showing intersection between lists of differentially expressed RNAs identified following pairwise comparison of each tumour group to NT control and pancreatitis groups GP1 and GP2. (E) Venn diagram showing intersection between lists of tumour specific differentially expressed RNAs from A – D. (F) Heatmap showing log2 fold-change relative to NT control for all upregulated RNAs in E. (G) Heatmap showing log2 fold-change relative to NT control for all downregulated RNAs in E. (H) Pie charts showing the percentage of different RNA species represented within the significantly up- and downregulated tumour specific RNAs in E.

### Tumour genotype impacts circulating miRNA biomarker expression

miRNAs are short non-coding oligonucleotides that can regulate gene expression, thereby impacting biological function. A number of miRNAs have been associated with cancer and their presence in biofluids, including blood, has led to interest in their diagnostic potential (22). To better understand the impact of tumour genotype on the utility of miRNAs in the blood as potential diagnostic biomarkers, total RNA was isolated from the blood of tumour bearing mice, NT and pancreatitis controls, and miRNA sequencing libraries prepared. Sequencing was performed on an Illumina NetSeq2000 and data analysed as described in materials and methods. PCA analysis across the six experimental groups identified limited intragroup variability with the exception of the pancreatitis group, which again appeared to split into two clusters (**Figure 4A**). To first understand whether the presence of a tumour, and its genotype, led to alterations in miRNA expression we performed pairwise comparisons between all four tumour genotypes and NT control to identify significantly differentially expressed miRNAs (padj ≤ 0.05, log2fc ≥ 1 or ≤ -1). This analysis identified 24 upregulated and 2 downregulated miRNAs in *Kras^G12D^ Trp53^-/-^*tumour bearing mice, 13 upregulated and 6 downregulated miRNAs in *Kras^G12D^ Trp53^-/-^ Cdkn2a^-/-^* tumour bearing mice, and 5 upregulated miRNAs in *Kras^G12D^ Trp53^-/-^ Smad4^-/-^*tumour bearing mice (**Figure 4B**). No differentially expressed miRNAs were identified from the blood of *Kras^G12D^ Trp53^-/-^ Cdkn2a^-/-^ Smad4^-/-^* tumour bearing mice. Cross-comparison of the miRNAs identified for the other three tumour groups showed that only 1 miRNA was common to all, and 9 were shared between *Kras^G12D^ Trp53^-/-^* and *Kras^G12D^ Trp53^-/-^ Cdkn2a^-/-^* groups.

**Figure 4.**
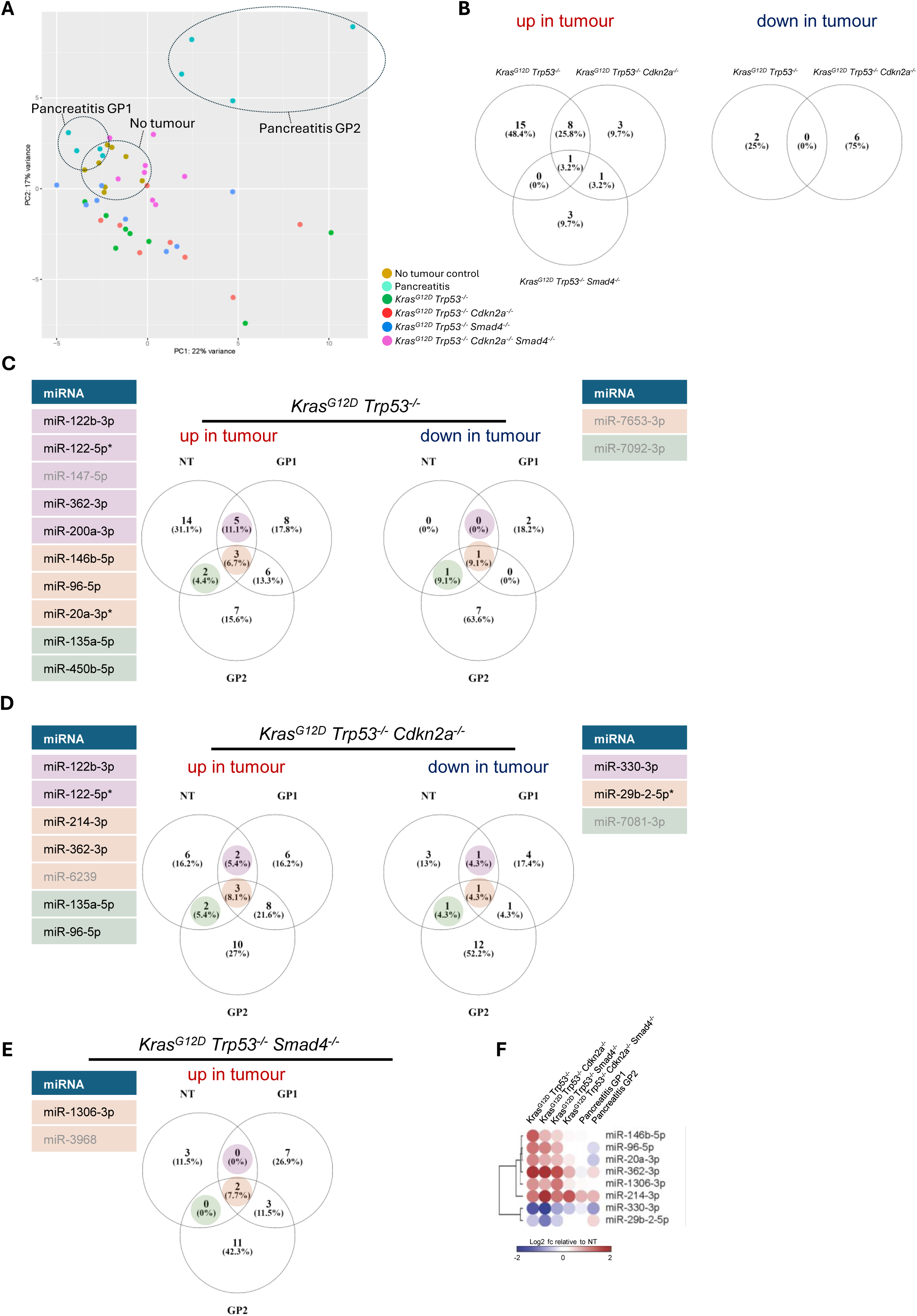
Tumour genotype impacts circulating miRNA biomarker expression. (A) Principal Component Analysis of miRNA sequencing data from biological replicates across all six experimental conditions. N = 8 biological replicates / group. (B) Venn diagram showing intersection between lists of differentially expressed miRNAs identified following pairwise comparison of each tumour group to NT control. (C) Venn diagram showing intersection between lists of differentially expressed miRNAs following pairwise comparison of miRNA expression from *Kras^G12D^ Trp53^-/-^*tumour bearing mice with NT controls, and pancreatitis groups GP1 and GP2. (D) Venn diagram showing intersection between lists of differentially expressed miRNAs following pairwise comparison of miRNA expression from *Kras^G12D^ Trp53^-/-^ Cdkn2a^-/-^* tumour bearing mice with NT controls, and pancreatitis groups GP1 and GP2. (E) Venn diagram showing intersection between lists of differentially expressed miRNAs following pairwise comparison of miRNA expression from *Kras^G12D^ Trp53^-/-^ Smad4^-/-^* tumour bearing mice with NT controls, and pancreatitis groups GP1 and GP2. Colours represent which segment of the Venn diagram the miRNAs belong. * = mouse specific miRNAs. (F) Heatmap showing log2 fold-change relative to NT control for all tumour specific miRNAs from C - E.

To further dissect the specificity of the differentially expressed miRNAs to the tumour setting, we next performed pairwise comparisons of each tumour genotype with both pancreatitis GP1 and GP2. Cross-comparison of differentially expressed miRNAs (padj ≤ 0.05, log2fc ≥ 1 or ≤ -1) relative to NT, GP1 and GP2 identified only 3 upregulated and 1 downregulated miRNA specific to the *Kras^G12D^ Trp53^-/-^*genotype (**Figure 4C**), 3 upregulated and 1 downregulated miRNA specific to the *Kras^G12D^ Trp53^-/-^ Cdkn2a^-/-^* genotype (**Figure 4D**), and 2 upregulated miRNAs specific to the *Kras^G12D^ Trp53^-/-^ Smad4^-/-^* genotype (**Figure 4E**). To better understand the level of specificity of each miRNA to a given tumour genotype we next visualised the log2fc relative to NT using a heatmap (**Figure 4F**). This analysis identified that the majority of ‘genotype-specific’ miRNAs were in fact upregulated across *Kras^G12D^ Trp53^-/-^*, *Kras^G12D^ Trp53^-/-^ Cdkn2a^-/-^*and *Kras^G12D^ Trp53^-/-^ Smad4^-/-^* groups, albeit to differing levels and therefore not reaching significance on all genotypes. However, most of these miRNAs were not upregulated in *Kras^G12D^ Trp53^-/-^ Cdkn2a^-/-^ Smad4^-/-^* tumour bearing mice, further confirming tumour genotype as a major barrier to identification of a pan-pancreatic cancer RNA-based biomarker signature.

### Tumour genotype dictates protein biomarker signatures in the plasma

In addition to RNA, proteins present in blood plasma represent a potentially valuable source of biomarkers to aid early detection and diagnosis efforts. Several plasma proteins have been identified as potential biomarkers for PDAC, but almost all have failed upon clinical validation (7, 8). To investigate the relationship between tumour genotype and protein expression in the plasma, we isolated plasma from blood 3-weeks post-implantation of either *Kras^G12D^ Trp53^-/^*^-^, *Kras^G12D^ Trp53^-/-^ Cdkn2a^-/-^*, *Kras^G12D^ Trp53^-/-^ Smad4^-/-^*or *Kras^G12D^ Trp53^-/-^ Cdkn2a^-/-^ Smad4^-/-^* cells into the pancreas of C57BL/6 mice, and analysed protein expression using mass spectrometry as described in materials and methods. PCA analysis based on proteins present in at least 50% of each experimental group identified limited intragroup variability (**Figure 5A**), with only one pancreatitis group being identified as opposed to the two distinct groups that had been observed when analysing RNAs. To gain insight into the relationship between plasma protein expression and tumour genotype, we first performed pairwise comparisons of each tumour group to NT controls to identify significant differentially expressed proteins (p-value ≤ 0.05, log2fc ≥ 0.58 or ≤ -0.58). This analysis identified 21 upregulated and 83 downregulated proteins for the *Kras^G12D^ Trp53^-/-^* genotype, 122 upregulated and 47 downregulated proteins for the *Kras^G12D^ Trp53^-/-^ Cdkn2a^-/-^*genotype, 16 upregulated and 53 downregulated proteins for the *Kras^G12D^ Trp53^-/-^ Smad4^-/-^* genotype, and 37 upregulated and 26 downregulated proteins for the *Kras^G12D^ Trp53^-/-^ Cdkn2a^-/-^ Smad4^-/-^* genotype (**Figure 5B**). Surprisingly, only 3 upregulated and 3 downregulated proteins were common to all four genotypes.

**Figure 5.**
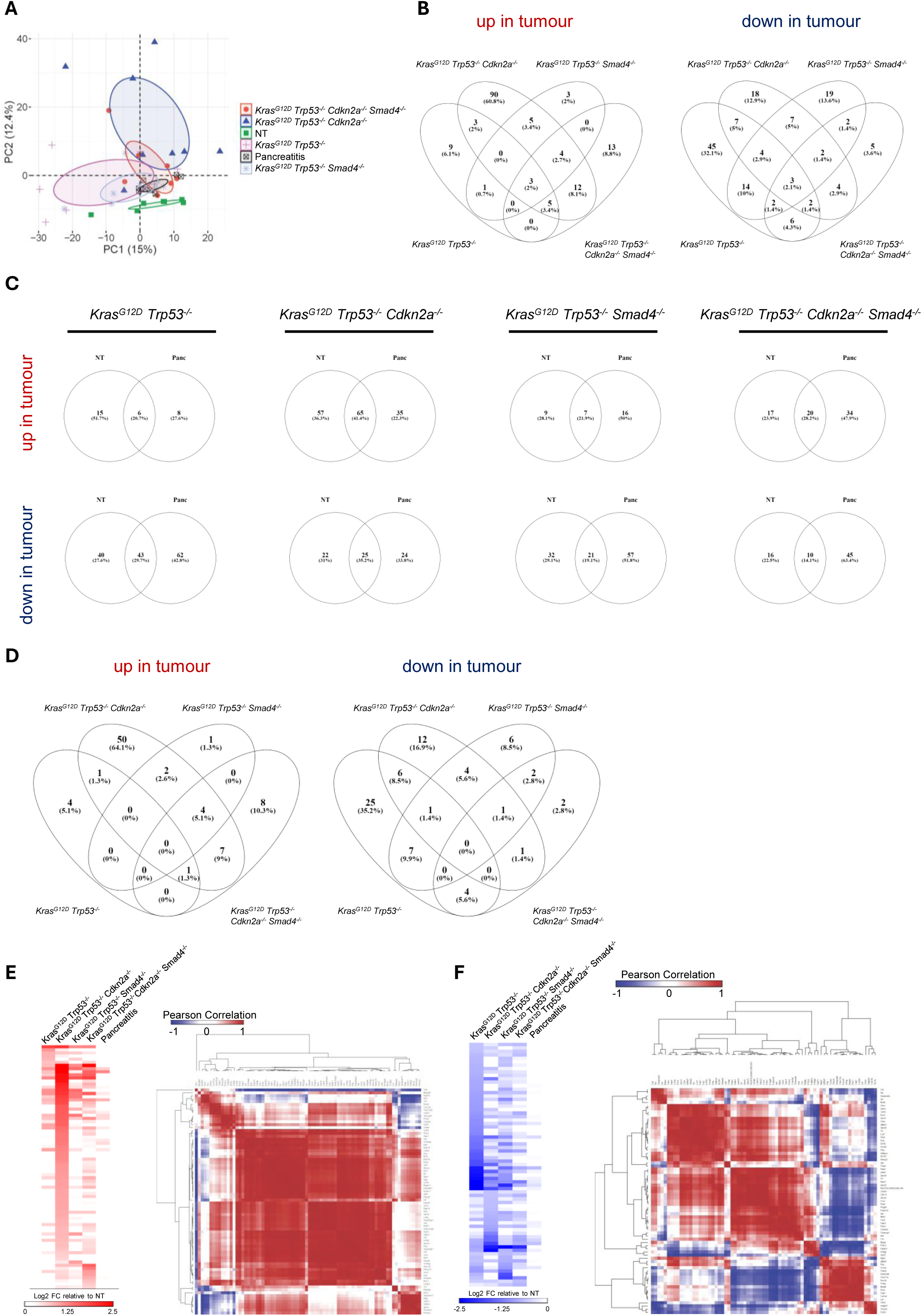
Tumour genotype dictates protein biomarker expression in the plasma. (A) Principal Component Analysis of plasma mass spectrometry proteomics data from biological replicates across all six experimental conditions. N = 8 biological replicates / group. (B) Venn diagram showing intersection between lists of differentially expressed proteins identified following pairwise comparison of each tumour group to NT control. (C) Venn diagram showing intersection between lists of differentially expressed proteins following pairwise comparison of each tumour group with NT and pancreatitis controls. (D) Venn diagrams showing intersection between lists of tumour specific differentially expressed proteins from C. (E) left – Heatmap of median centred log2 expression values for significantly upregulated tumour specific proteins in D, right – Pearson correlation analysis and hierarchical clustering of tumour specific upregulated proteins in D. (F) left – Heatmap of median centred log2 expression values for significantly downregulated tumour specific proteins in D, right – Pearson correlation analysis and hierarchical clustering of tumour specific downregulated proteins in D.

To further determine whether the proteins identified were specific to the tumour setting or associated with more generalised pancreatic inflammation, we next performed pairwise comparisons of all four tumour groups to pancreatitis controls. Cross-comparison of significant differentially expressed proteins (p-value ≤ 0.05, log2fc ≥ 0.58 or ≤ -0.58) to both NT and pancreatitis controls identified that a substantial proportion of proteins regulated relative to NT controls were associated with more generalised inflammation and would therefore not provide the specificity to discriminate between PDAC and inflammation of the pancreas. This analysis resulted in identification of 6 upregulated and 43 downregulated proteins for the *Kras^G12D^ Trp53^-/-^* genotype, 65 upregulated and 25 downregulated proteins for the *Kras^G12D^ Trp53^-/-^ Cdkn2a^-/-^* genotype, 7 upregulated and 21 downregulated proteins for the *Kras^G12D^ Trp53^-/-^ Smad4^-/-^* genotype, and 20 upregulated and 10 downregulated proteins for the *Kras^G12D^ Trp53^-^*

*^/-^ Cdkn2a^-/-^ Smad4^-/-^* genotype that represented potential tumour-specific biomarkers (**Figure 5C**). Cross-comparison of these refined protein lists showed that no tumour-specific differentially expressed proteins were common across all four genotypes when pancreatitis was also taken into consideration (**Figure 5D**). Furthermore, heatmap visualisation of log2fc relative to NT controls further supported the specificity of plasma protein signatures to a given tumour genotype or subgroup of genotypes (**Figures 5E** and **5F**). Notably, the plasma of hosts bearing *Kras^G12D^ Trp53^-/-^ Cdkn2a^-/-^* tumours had by far the most upregulated proteins when compared with NT and pancreatitis controls, with cell type and GO biological process overrepresentation analysis (**Supplementary Figure 3**) suggesting that these may be the consequence of altered immune cell abundance, in particular neutrophils, or immune cell phenotype. Collectively, these findings imply that tumour genotype is likely to be a major confounding factor in the identification of a pan-PDAC biomarker signature, supporting the need to move towards genotype-specific biomarkers when using plasma proteins.

A number of multi-protein biomarker panels have been proposed as potential candidates to support in the earlier detection of PDAC (9) based on discovery efforts using human samples. However, to-date, none of these have been successfully validated and adopted into clinical practice. To gain a better understanding of whether tumour genetic background may impact the diagnostic performance of these panels, we extracted a list of proteins from our plasma MS proteomics dataset that were represented across four different previously published human panels (23–26). A heatmap of median centred log2 expression values across all four tumour genotypes, NT controls and pancreatitis controls identified genotype-specific alterations in protein expression (**Figure 6A**). Pearson correlation analysis identified good correlations between different sets of proteins (**Figure 6B**). However, in general these did not correspond to the composition of the previously reported panels (**Figure 6C**). Therefore, in all cases tested, our data implies that tumour genotype would degrade diagnostic performance, supporting the need to understand the impact of tumour genotype on biomarker expression when designing multi-marker diagnostic panels.

**Figure 6.**
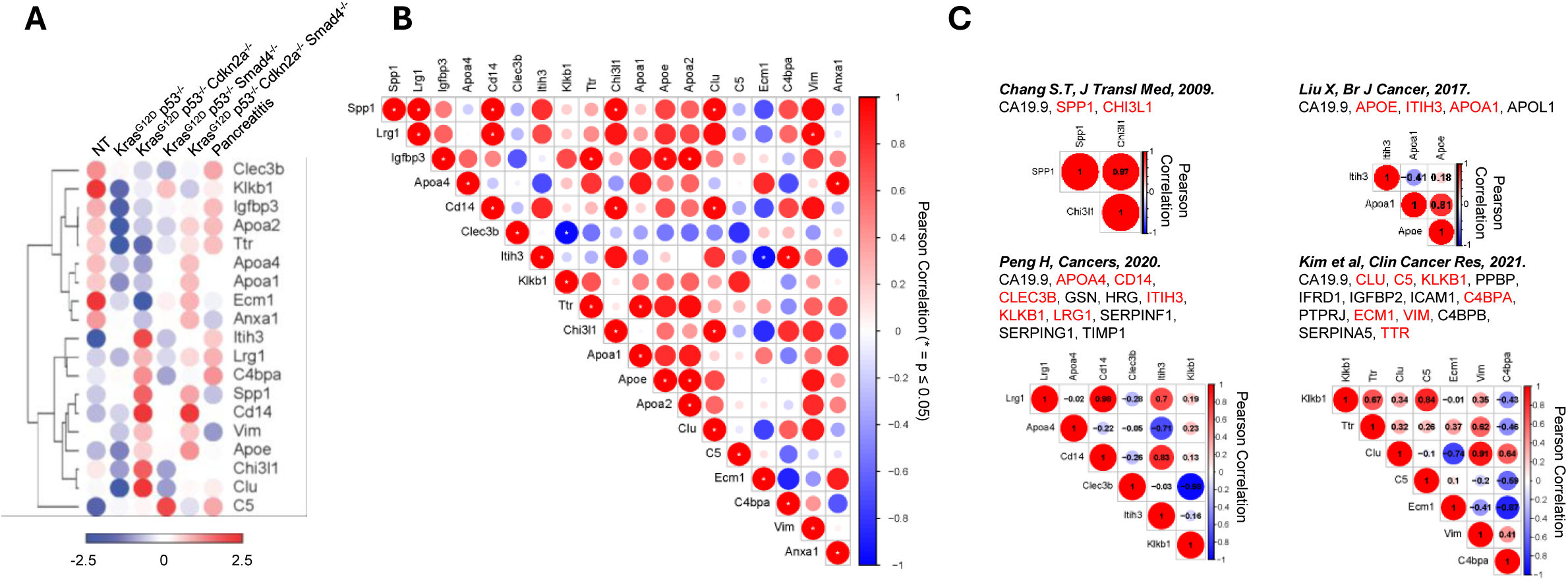
Tumour genotype degrades performance of previously published human diagnostic serum protein biomarker panels. (A) Hierarchical clustering of median centred log2 expression values for a range of previously reported PDAC protein biomarkers across all experimental conditions. (B) Pearson correlation analysis for a range of previously reported PDAC protein biomarkers using log2 expression values from all tumour groups. (C) Pearson correlation analysis based on four previously published diagnostic plasma / serum protein biomarker panels using log2 expression values from all tumour groups. N = 32 tumour plasma samples. Proteins highlighted in red are present within the plasma MS-proteomic dataset generated as part of this study.

## DISCUSSION

Here, we use a novel genetically defined model system to directly address whether genetic heterogeneity inherent to human PDAC likely represents a major confounding factor in ongoing efforts to identify biomarkers to support earlier detection of this highly lethal disease. Using multi-omics analysis of blood, we find that common genetic lesions i.e. loss of *Cdkn2a* and or *Smad4*, profoundly alter the systemic pool of available RNA and protein biomarkers, resulting in few pan-genotype but many more genotype-specific biomarkers. These data provide a plausible explanation for failure of current candidate biomarkers and strongly support the need to incorporate genomic information into current and future clinical biomarker studies.

The majority of biomarker discovery efforts focus on the use of human biospecimens. However, many studies do not focus on early-stage patients and have relatively small sample sizes (9), the latter resulting in low statistical power and the inability to probe relationships between defined patient variables and biomarker performance. Given the relatively low incidence of PDAC, together with the inherent genetic and molecular heterogeneity, and patient variables including sex, ethnicity, diet, lifestyle etc, larger cohorts sourced from multiple centres will almost certainly be required to adequately assess biomarkers. This introduces additional challenges related to sample processing, storage and standardisation of methods used for biomarker detection, highlighting the potential limitations of human studies for determining direct cause and effect relationships. While one can argue the merits of using mouse models for bona-fide biomarker discovery (27), by virtue of their defined genetics, limited interpatient heterogeneity, and highly standardized lifestyle, they make ideal model systems for establishing direct cause and effect relationships, thereby helping to shape the design of clinical studies.

It is noteworthy that multiple candidate biomarkers that have been tested for diagnostic power in human clinical cohorts were present within the datasets generated here. For example, osteopontin (SPP1, OPN), which has been tested as a plasma biomarker both alone and in combination with additional proteins including CA19.9 and CHI3L1 (23), showed specific enrichment only in the plasma of mice bearing tumours harbouring Cdkn2a loss. Therefore, osteopontin may still represent a valuable biomarker for the detection of PDAC, but in the context of a larger multi-marker panel that has the potential to inform on the underlying genetics of a patient’s tumour. Several multi-marker panels have been proposed previously but again have failed to reach clinical implementation (24–26). Pearson correlation analysis of such panels based on the data generated here, strongly suggests that tumour genotype would degrade the diagnostic accuracy of all panels tested, again supporting the notion that understanding the relationship between tumour genotype and biomarker expression could improve the design and implementation of large multi-marker diagnostic panels.

This study has focussed on understanding the impact of CDKN2A and SMAD4 loss-of-function in the context of a KRAS^G12D^ mutant allele. KRAS mutation occurs in >90% of human PDAC, with the KRAS^G12D^ mutation present in approximately 50% of cases. Therefore, while KRAS^G12D^ is the most common allele, alternative KRAS variants including KRAS^G12V^ (∼30%) and KRAS^G12R^ (∼15%) are also common in PDAC (10). Growing evidence supports the conclusion that different KRAS mutant proteins do not drive effector signalling equally (28–30), suggesting that the KRAS allele present within a tumour may further impact the pool of biomarkers expressed in the blood. Based on our findings, we propose that a more in-depth understanding of the extent to which genetics influences biomarker expression, incorporating all major KRAS variants and additional driver mutations, will be critical to improving the chances of developing clinically implementable diagnostic panels to aid earlier detection of pancreatic cancer. The approach we report here i.e. the use of CRISPR-Cas9 genome editing to generate malignant cells from genetically pristine primary epithelial cells of the pancreas, represents an economically viable, scalable and rapid solution to addressing this complex question, thereby enabling a clear understanding of the extent to which genomic information will be important in future clinical biomarker discovery efforts.

While this study has been carried out in the context of pancreatic cancer, our findings may be broadly applicable across cancers. Loss or mutation of CDKN2A is common in a range of cancer types, including melanoma, mesothelioma, head and neck, bladder, gastrointestinal stromal tumours, non-small cell lung and esophagogastric cancers (31). Therefore, it is possible that the broad reprogramming of blood biomarker expression that we observe here to be driven by loss of CDKN2A could also impact biomarker identification and validation efforts for many cancers. It is likely that this may also be the case for cancers with loss of, or mutation in, SMAD4. While this is most common in pancreatic cancer, it also occurs at lower frequency in many other cancer types including colorectal, esophagogastric, stomach, non-small cell lung, endometrial, bladder, cholangiocarcinoma and ovarian cancers (32). Therefore, while our observations that loss of SMAD4 severely limits the pool of available biomarkers may have the greatest impact on pancreatic cancer diagnosis, it remains possible that it also presents an unappreciated roadblock in the identification and validation of RNA blood-based biomarkers for early detection more generally. While this study has focussed on loss of CDKN2A and SMAD4, genetic heterogeneity is common to almost all cancers (33) and our findings here suggest that incorporating this information into future biomarker discovery efforts will be important.

## MATERIALS AND METHODS

### Isolation and CRISPR-Cas9 gene editing of primary acinar cells

Female C57BL/6 mice were euthanised by cervical dislocation and the pancreas removed and washed 2x with HBSS (Life Technologies). The pancreas was subsequently cut into small pieces using a scalpel and digested in 10 ml of HBSS containing 10 mM HEPES (Life Technologies), 200 U/ml collagenase IA (Sigma-Alrich), and 0.25 mg/ml of trypsin inhibitor (Sigma-Aldrich) at 37°C for 30 minutes, shaking at 200 rpm. Digestion was stopped by addition of 10 ml of cold wash solution (1x HBSS supplemented with 5% Foetal Bovine Serum (FBS, Life Technologies)). The resulting cell suspension was centrifuged 2 minutes at 4°C, 450 xg and the supernatant discarded. The cell pellet was resuspended in 10 ml of wash buffer and this process repeated two more times. The cell pellet was then resuspended in 7 ml of Waymouth’s medium (Life Technologies) containing 2.5% FBS, 1% penicillin-streptomycin (Life Technologies), 0.25 mg/ml of trypsin inhibitor (Sigma), and 25 ng/ml of recombinant human Epidermal Growth Factor (EGF, Gibco). Cell suspension filtered through a 100 µm filter, and acini cultured in a 6-well dish (2 ml per well) at 37 °C with 5% CO2. After 24 hours, acini in suspension were transferred to a collagen coated 60 mm dish (Corning Biocoat) and culture media changed every 48 hours. Acinar cells cultured for 4 days and subjected to CRISPR/Cas9 editing as follows.

Reagents to generate RNP complexes were ordered through IDT’s Alt-R platform. RNP complexes were prepared as per Dewari et al. (2018) (34). Both tracrRNA and crRNA (**Table 1**) were resuspended in nuclease-free duplex buffer (IDT) at 100 µM. For one reaction of up to 500,000 cells, 1.1 µL of each the crRNA and tracrRNA were mixed in a sterile 0.2 mL PCR tube. crRNA and tracrRNA were annealed to form guide RNA duplex by heating to 95°C for 5 minutes and allowed to cool to room temperature for 30 minutes before being plunged into ice. 1 µL of 61 µM stock of Alt-R Cas9 V3 HiFi was added to the guide RNA duplex and incubated at room temperature for 10 minutes to allow for RNP formation. For increased probability of inducing a double-strand break (DSB) within the target gene, RNP complexes with two separate crRNA sequences were generated for TRP53 KO. Single stranded oligodeoxynucleotide (ssODN) repair template for KRAS^G12D^ incorporation via HDR (table 1) was resuspended in nuclease-free duplex buffer at 30 µM.

**Table 1.**
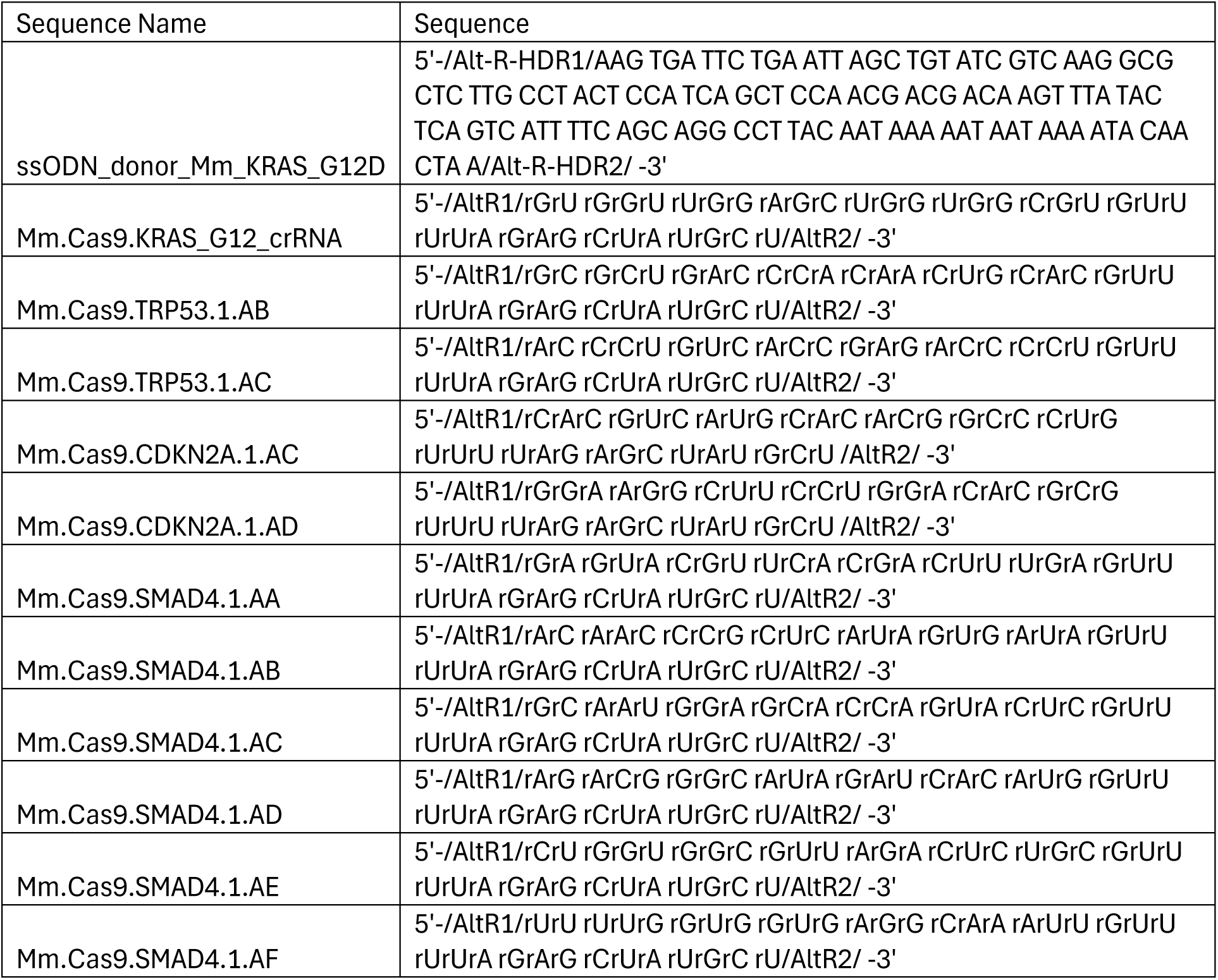

300,000 to 500,000 acinar cells were harvested using TrpLE (Life Technologies), resuspended in 20 µL SE cell line buffer (Lonza SE Cell Line 4D-Nucleofector X Kit S) and 1 µL nucleofection enhancer was added. 3.2 uL of each RNP complex and 1 µL ssODN were added to reaction mixture and entire volume (∼30 µL) transferred to one well in a 16-well nucleocuvette (Lonza). Cells were electroporated in an Amaxa 4D nucleofector X unit (Lonza) with program EO-100, then transferred into a 60 mm collagen I coated dish containing 4 mL pre-warmed Waymouth’s medium (supplemented as above). For HDR based KI, cells cultured in medium supplemented with 30 µM HDR enhancer (IDT) for 24 hours and cells incubated at 32°C /5% CO2 (cold-shock) for 48 hours as per Skarnes et al. (2019) (35).

To isolate clones expressing KRAS^G12D^ and negative for expression of p53, a confluent 100 mm dish was harvested using cell dissociation buffer (Millipore) at 37 °C for 10 minutes, passed through a 70 µm filter and single cells seeded into each well of a 96-well plate using a Beckman Coulter CytoFLEX SRT flow cytometer. Clones were expanded and tested for expression of KRAS^G12D^ and p53 using western blotting. Following identification of a clone, *Cdkn2a* and *Smad4* expression was knocked out using the same CRISPR-Cas9 approach as that detailed above. For *Smad4* KO, RNP complexes were generated with six unique crRNA for improved knock-out efficiency. No ‘cold shock’ and subsequent single cell cloning was performed for gene knockout.

All cell lines were cultured at 37°C with 5% CO2 in high-glucose Dulbecco’s Minimum Essential Medium (Sigma) supplemented with 10% fetal bovine serum (Life Technologies). Cell lines were routinely tested for mycoplasma every 2–3 months in-house and were mycoplasma-negative. Cell lines are cultured for no more than 3 months following thawing.

### Western blotting

Cell lysates were prepared using RIPA buffer (50 mmol/L Tris-HCl, pH 7.6, 150 mmol/L sodium chloride, 1% Triton X-100, 0.5% deoxycholate, 0.1% SDS) supplemented with phosphatase inhibitor cocktail (Roche) and protease inhibitor cocktail (Roche). 10 – 20 μg of protein, as measured by Micro BCA Protein Assay kit (Pierce), was supplemented with SDS sample buffer, separated by SDS–PAGE, transferred to nitrocellulose and immunoblotted with anti-Kras^G12D^ (1/1000, Cell Signalling Technologies, #14429), p53 (1/1000, Cell Signalling Technologies, #2524), p16^INK4A^ (1/1000, Cell Signalling Technologies, #29271), p19^ARF^ (1/1000, Cell Signalling Technologies, #77184), Smad4 (1/1000, Cell Signalling Technologies, #46535), Keratin-19 (1/1000, Cell Signalling Technologies, #3984) and α-tubulin (1/2000, Cell Signalling Technologies, #3873) antibodies. Fluorescent detection was carried out following incubation with DyLight 680/800-conjugated secondary antibodies using a LI-COR Odyssey CLx scanner (LI-COR Biosciences).

#### RPPA

Cell lysates were prepared using RIPA buffer (50 mmol/L Tris-HCl, pH 7.6, 150 mmol/L sodium chloride, 1% Triton X-100, 0.5% deoxycholate, 0.1% SDS) supplemented with phosphatase inhibitor cocktail (Roche) and protease inhibitor cocktail (Roche). Protein concentration was determined using a Micro BCA Protein Assay kit (Pierce). Samples were prepared for printing by addition of 4x sample buffer followed by heat denaturation at 95°C for 5 minutes. All samples were diluted to a range of concentrations: 1.5 mg/ml, 0.75 mg/ml, 0.375 mg/ml and 0.1875 mg/ml using PBS containing 10% glycerol. All four concentrations of each sample were spotted onto single pad ONCYTE SuperNOVA nitrocellulose slides at a spot-to-spot distance of 500 µm using a Quanterix 2470 Arrayer platform equipped with 185 µm pins, with one round of deposition. Following spotting, slides were incubated with antigen retrieval solution (1x Reblot strong) for 10 minutes before being clamped into a hybridisation cassette to divide the slide into subarrays. Blocking buffer (Superblock T20) was then added to each subarray and incubated for 10 minutes. Blocking buffer was then removed and primary antibody diluted in blocking buffer added for 60 minutes at room temperature. Primary antibody was then removed, each subarray washed with PBS-T, followed by a further incubation with blocking buffer and another wash with PBS-T. Secondary antibody (Dylight-800 conjugated, species matched, 1:2500 dilution in Superblock T20) was then added to each subarray and incubated for a further 30 minutes at room temperature in the dark. Slides were then washed with PBS-T before being dried and imaged using an Innopsys Innoscan 710 microarray scanner. Non-specific background fluorescence was determined by omitting the addition of primary antibody. The average fluorescent intensity of each spot in the resulting images was determined using Mapix software (Innopsys) with the spot diameter set to 270 µm. A linear fit was applied to the 4-point dilution series to ensure that detection was within the dynamic range of the assay and median values from the 4-point dilution series calculated for analytes with an R^2^ ≥ 0.8. Relative fluorescent intensity values were z-transformed and hierarchically clustered (one minus Pearson correlation, average linkage) using the Morpheus software package (https://software.broadinstitute.org/morpheus).

### Orthotopic implantation of cancer cells into the pancreas

Female C57BL/6 mice (Envigo) were supplied as age-matched, 5-week-old females and isolated for 1 week after delivery. Mice were anaesthetised with inhalational isoflurane anaesthetic in oxygen, and administered perioperative analgesia (buprenorphine (Vetergesic, 0.1 mg/kg s.c.)). Post-surgery mice were also administered carprofen via their drinking water for 48 hours. Cell lines were propagated to subconfluency to ensure they were in their exponential growth phase. Once detached from the flask and washed with PBS, cells were suspended in growth factor-reduced matrigel (Scientific Laboratory Supplies) at a concentration of 1×10^5^ cells in 10 µL. Using aseptic technique, a 3 mm skin incision was made in the left lateral flank and lateral abdominal muscles in order to visualise the pancreas. 10 µL of matrigel containing cells was injected into the pancreas in a sterile manner. The abdominal wall was closed with Polyglactin 910 (Vicryl, 2M, Henryschein) using a single cruciate suture. Skin was closed with skin clips. Mice were monitored in a heat box set to 37°C post-surgery for 1 hour. Mice were closely monitored daily with twice-weekly weight checks following implantation. If any single terminal symptom caused by pancreatic tumour growth, including weight loss equal to or exceeding 10% of the starting weight, signs of abdominal pain or abdominal distension became apparent, the animal was humanely euthanised. After 21 days, the animals were culled (cervical dislocation) and the pancreatic tumours were harvested for analysis. Tumour weights were recorded and agreed by two observers. In addition, 0.5 ml of blood was obtained by cardiac puncture. All experiments had University of Edinburgh ethical approval and were carried out in accordance with the United Kingdom Animal Scientific Procedures Act (1986).

### Cerulein-induced pancreatitis

Female C57BL/6 mice (6-8 weeks old) were fasted for 15 hours. To induce acute pancreatitis, cerulein (80 μg/kg) (MedChemExpress) was administered via intraperitoneal injection a total of 6 times, spaced at 1-hour intervals. Mice were also administered buprenorphine (Vetergesic, 0.1 mg/kg) subcutaneously as an analgesic 30 minutes before first cerulein dose. Mice were sacrificed (exposure to CO2 gas in rising concentration) two days after first cerulein dose, and blood was harvested via cardiac puncture.

### RNA Isolation

30 mg of tumour tissue, stored in RNALater at -80°C, was used to isolate RNA using a Qiagen RNeasy Mini kit, including DNase digestion, as per the manufacturer’s instructions.

Whole blood RNA extracts were obtained using Mouse RiboPure Blood RNA isolation kit (Ambion Inc), following manufacturer’s instructions.

Total RNA was assessed on the Fragment Analyser Automated Capillary Electrophoresis System (Agilent Technologies Inc, #5300) with the Standard Sensitivity RNA Analysis Kit (#DNF-471-0500) for quality and integrity of total RNA, then quantified using the Qubit 2.0 Fluorometer (Thermo Fisher Scientific Inc, #Q32866) and the Qubit RNA broad range assay kit (# Q10210). DNA contamination was quantified using the Qubit dsDNA HS assay kit (#Q32854).

### Total RNA sequencing

Libraries were prepared from total-RNA samples using the NEBNext Ultra 2 Directional RNA library prep kit for Illumina (#E7760) and the NEBNext globin and rRNA Depletion kit (Human/Mouse/Rat) (#E7750) according to the provided protocol. 250ng of total-RNA was added to the globin and ribosomal RNA (rRNA) depletion reaction using the NEBNext globin and rRNA depletion kit (Human/mouse/rat). Globin / rRNA-depleted RNA was then DNase treated and purified using Agencourt RNAClean XP beads (Beckman Coulter Inc, #66514). RNA was then fragmented using random primers before undergoing first strand and second strand synthesis to create cDNA. Fragmentation time was dependent upon the quality of each total RNA sample. cDNA was end-repaired before ligation of sequencing adapters, and adapter-ligated libraries were enriched by 11 cycles of PCR using NEBNext Multiplex Oligos for Illumina (96 Unique Dual Index Primer Pairs) (NEB, #E6440). Final libraries had an average peak size of 286bp.

Libraries were quantified by fluorometry using the Qubit dsDNA HS assay and assessed for quality and fragment size using the Agilent Bioanalyser with the DNA HS Kit (#5067-4626). Fragment size and quantity measurements were used to calculate molarity for sequencing. Sequencing (2×100) was performed on the NextSeq 2000 platform (Illumina Inc, #20038897) using NextSeq 2000 P3 Reagents (200 Cycles) (#20040560). Libraries were combined in 1 equimolar pool of 48 based on Qubit and Bioanalyser assay results and run over two P3 flow cells. Coverage was broadly even across the flow cells and the majority of libraries generated ≥50M PE reads (Min: 41.2M, Max: 70.3M, Mean: 53.8M).

### miRNA sequencing

Sequencing libraries were prepared from 100ng of each total RNA sample using the QIAseq miRNA library kit (QIAGEN, #331502) and the QIAseq miRNA NGS 48 Index IL kit (#331595) according to the provided protocol. Mature miRNAs possess both a 3’ hydroxyl group and a 5’ phosphate. A pre-adenylated DNA adapter was ligated to the 3’ ends of all miRNAs and then an RNA adapter was ligated to the 5’ end of mature miRNAs. After both ligations, the ligated miRNAs were reverse transcribed into cDNA. The reverse transcription primer contained an integrated UMI and bound to a region of the 3’ adapter where it facilitates conversion of the ligated miRNAs into cDNA while assigning a UMI to every miRNA molecule. During this step a universal sequence was also added that is recognized by the sample indexing primers during library amplification. Libraries were purified using QIASeq miRNA NGS Beads (QMN beads). Purified libraries were then amplified for 16 cycles of PCR with 6bp single index sequences to allow multiplexing on a single sequencing run. Amplified indexed libraries were purified and size selected with QMN beads to enrich for library fragments from miRNA. Libraries were quantified by fluorometry using the Qubit dsDNA HS assay and assessed for quality and fragment size using the Agilent Bioanalyser with the DNA HS Kit (#5067-4626). Molarity for sequencing was calculated using the quantification and fragment size data. Sequencing (1×75) was performed on the NextSeq 2000 platform (Illumina Inc, #20038897) using NextSeq 1000/2000 P2 Reagents (100 Cycles) v3 (#20046811). PhiX Control v3 library (Illumina, # FC-110-3001) was spiked in at a concentration of 1% to enable troubleshooting in the event of run failure. Sequencing was single-end 1×75 to allow coverage of the UMI sequence and use in subsequent analyses. Coverage was broadly even and all but one library generated ≥8M reads (Min: 6.9M, Max: 13.7M, Mean: 10.4M).

### RNA and miRNA sequencing analysis

*Sequence alignment*: Data was processed using nf-core/rnaseq v3.14.0 (doi: https://doi.org/10.5281/zenodo.1400710) of the nf-core collection of workflows (36) for RNAseq and for miRNAs, nf-core/smrnaseq v2.3.1 (doi: https://zenodo.org/badge/latestdoi/140590861). The pipelines were executed with Nextflow v23.10.1 (37). Reads were aligned to the GRCm38 mouse genome.

*Differential expression analysis*: The DESeq2 package (version 1.42.1) in R (version 4.3.3) was used to perform differential gene expression analysis between various genotypes (38). Gene counts were obtained from the nf-core/rnaseq pipeline output and imported into DESeq2 using the DESeqDataSetFromMatrix method. Counts were normalised using the variance stabilising transformation (VST) and principal component analysis (PCA) performed to visualise sample clustering. Deseq2 fitted a negative binomial generalised linear model with genotype as the main factor and used the Wald test to identify differentially expressed genes. Genes were annotated using the biomaRt package (version 2.58.2) and the org.Mm.eg.db package (version 3.18.0). Hierarchical clustering (one minus Pearson correlation, average linkage) and heatmaps were generated using the Morpheus software package (https://software.broadinstitute.org/morpheus). Venn diagrams were generated using Venny2.0. Cell type and gene ontology biological process over-representation analysis was performed using the WEB-based Gene SeT AnaLysis Toolkit (webgestalk.org) with a reference set of the protein-coding genome and an FDR of ≤ 0.05. Enrichment bubble plots were generated using SRplot (39).

### Mass Spectrometry Proteomics analyses

Blood (0.5ml) was collected by cardiac puncture into EDTA-treated tubes. Cells were removed from plasma by centrifugation for 15 minutes at 2,000 x g using a refrigerated centrifuge. 2ul of the resulting supernatant (plasma) was trypsin digested using the PAC digest protocol (40) on a KingFisher Flex (Thermo Fisher) and analysed using a Data-independent-acquisition (DIA) workflow. Briefly, 2ug of peptides was analysed using a Fusion Lumos mass spectrometer coupled to an RSLCnano HPLC system as described in (41). Data from up to 8 biological replicates from each experimental group were analysed using DIA-NN 1.9 software (42) in tDIA mode and searched against the mouse Uniprot database.

Mass spectrometry proteomics data was analysed using Perseus software (version 2.1.3.0). LFQ values were log2 transformed and the protein list filtered based on a minimum requirement of 50% valid values in each group. Missing values in the resulting data matrix were then imputed based on a normal distribution and pairwise comparisons performed based on a students t-test, p-value threshold of ≤ 0.05. Principal Component Analysis was performed using SRplot (39) based on the filtered expression matrices. Venn diagrams were generated using Venny2.0. Hierarchical clustering (one minus Pearson correlation, average linkage) and heatmap visualisation was performed using Morpheus software (https://software.broadinstitute.org/morpheus). Pearson correlation coefficients were calculated using SRplot (39) and hierarchical clustering (precomputed similarity matrix, average linkage) and heatmap visualization performed using the Morpheus software package. Cell type and gene ontology biological process over-representation analysis was performed using the WEB-based Gene SeT AnaLysis Toolkit (webgestalk.org) with a reference set of the protein-coding genome and an FDR of ≤ 0.05. Enrichment bar plots and bubble plots were generated using SRplot (39).

### Statistical analyses

Statistical analysis was carried out using GraphPad Prism10 for Windows (GraphPad Software). All error bars on graphs represent standard error of the mean (sem). Statistical tests are detailed in the figure legends. n numbers provided for each experiment in the figure legends represent biological replicates.

## FIGURE LEGENDS

**Supplementary Figure 1.**
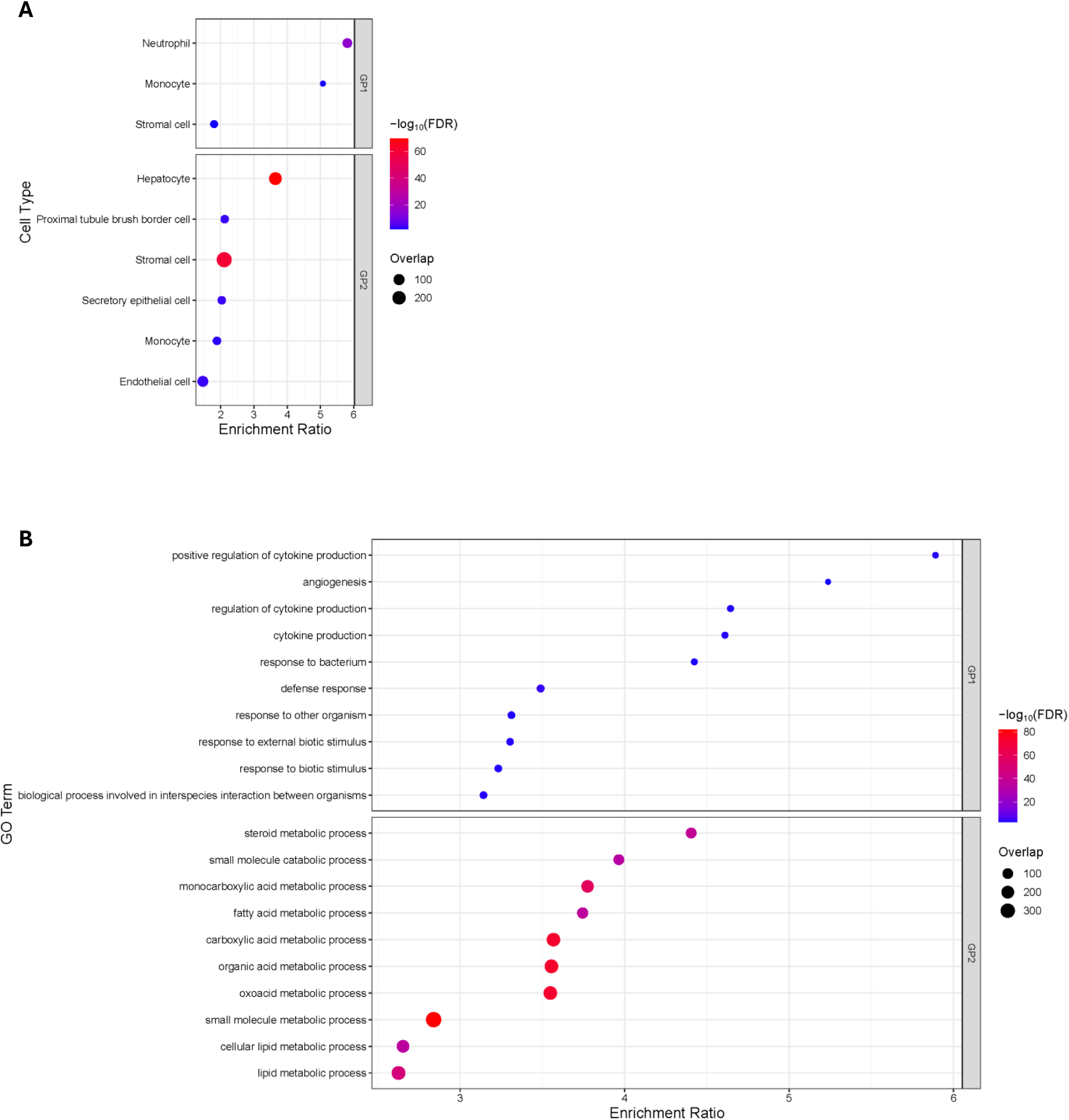
Comparison of pancreatitis groups GP1 and GP2. (A) Cell type overrepresentation analysis based on genes significantly upregulated in pancreatitis GP1 and GP2 following pairwise comparison with NT controls. (B) Gene ontology biological process overrepresentation analysis based on genes significantly upregulated in pancreatitis GP1 and GP2 following pairwise comparison with NT controls.

**Supplementary Figure 2.**
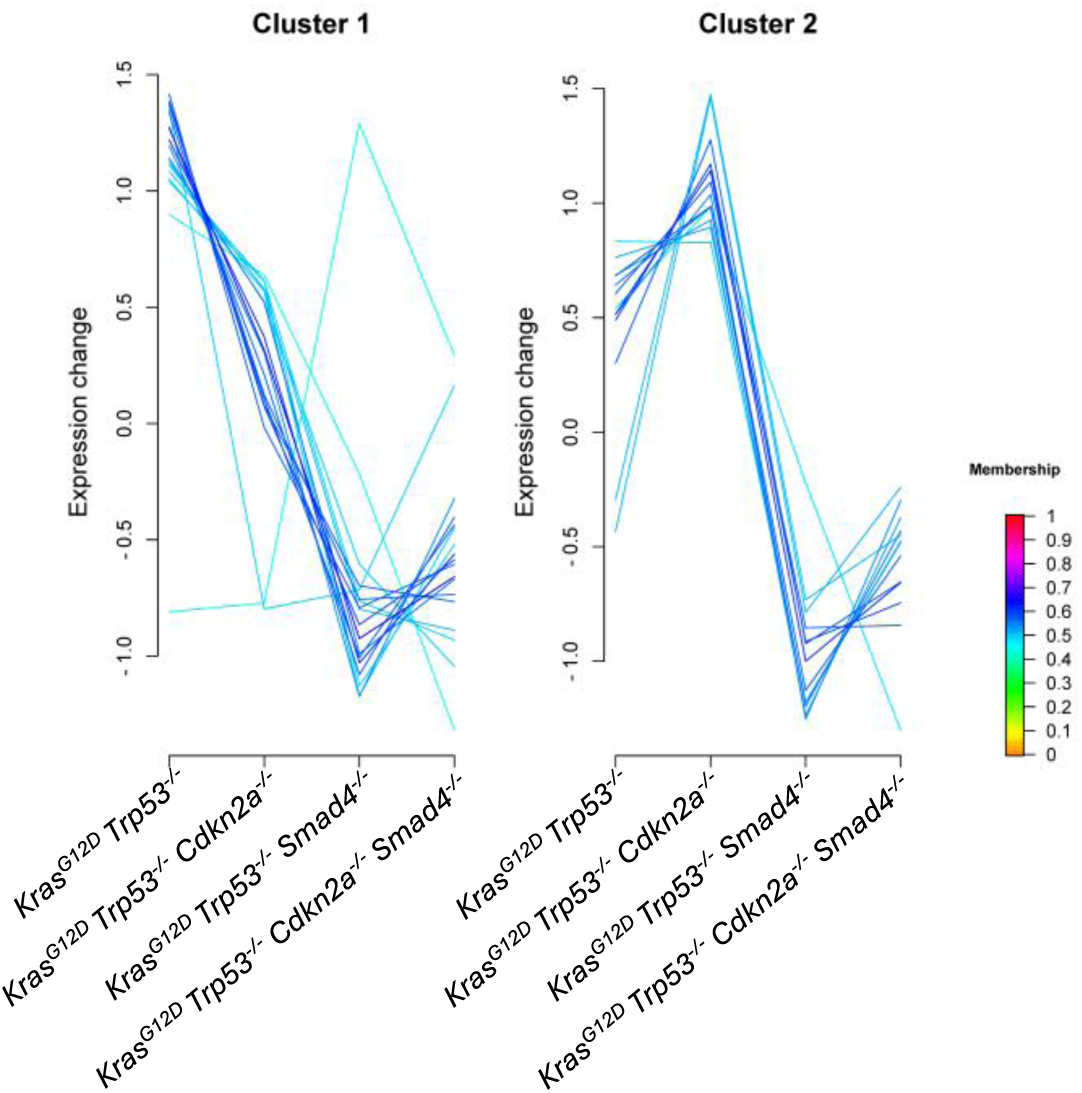
*Smad4* loss alone or in combination with *Cdkn2a* negatively impacts the log2 fold-change of 31 RNAs common to all four tumour genotypes. Line graphs showing the log2 fold-change of 31 RNAs from figure 2D that were upregulated across all four tumour genotypes following pairwise comparison with NT controls. Cluster 1 and cluster 2 represent groups of RNAs with similar expression profiles.

**Supplementary Figure 3.**
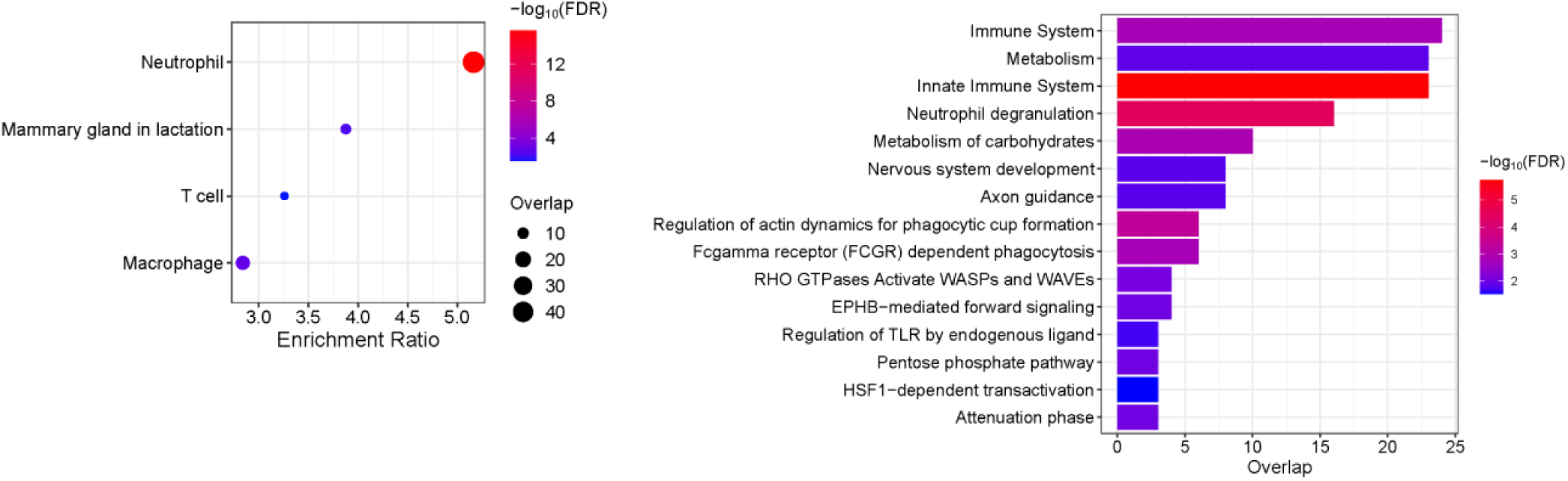
Cell type and Gene Ontology Biological Process enrichment in *Kras^G12D^ Trp53^-/-^ Cdkn2a^-/-^*plasma. (A) Cell type overrepresentation analysis based on proteins specifically upregulated in *Kras^G12D^ Trp53^-/-^ Cdkn2a^-/-^*plasma. (B) Gene ontology biological process overrepresentation analysis based on proteins upregulated specifically in *Kras^G12D^ Trp53^-/-^ Cdkn2a^-/-^* plasma.

## ACKNOWLEDGMENTS

We would like to thank the University of Edinburgh Central Biological Services animal technicians at WGH for their assistance with animal research studies, the Genetics Core at the Clinical Research Facility Edinburgh for their assistance with RNA library preparation and sequencing and the Host and Tumour Profiling Unit at the Institute of Genetics and Cancer for their assistance with RPPA studies.

## Author Contributions

MC and AS devised and oversaw the project. MC and AS designed the experiments with contributions from JPM and AVK. MC, AS, DL, AVK and CF performed the experiments. PG performed initial bioinformatics analysis of RNAseq and miRNAseq data. AS and MC performed additional bioinformatic analysis of RNAseq and miRNAseq data. AS and CF performed bioinformatic analysis of plasma proteomic data. MC and AS wrote the manuscript with contributions from JPM and AVK; all authors commented on and approved the final version. MC and AS are responsible for the overall content as guarantor.

## Data Availability

RNA sequencing data is available to download from NCBI GEO with accession numbers GSE302170 and GSE302035. Mass spectrometry data is available via ProteomeXchange with identifier PXD065456.

